# Repurposing of the tamoxifen metabolites to treat methicillin-resistant *Staphylococcus epidermidis* and vancomycin-resistant *Enterococcus faecalis* infections

**DOI:** 10.1101/2021.05.20.445078

**Authors:** Andrea Miró-Canturri, Andrea Vila-Domínguez, Rafael Ayerbe-Algaba, J Pachón, Manuel E. Jiménez-Mejías, Younes Smani

**Affiliations:** Clinical Unit of Infectious Diseases, Microbiology and Preventive Medicine, Virgen del Rocío University Hospital, Seville, Spain; Institute of Biomedicine of Seville (IBiS), Virgen del Rocío University Hospital /CSIC/University of Seville, Seville, Spain; Department of Medicine, University of Seville, Seville, Spain

## Abstract

Repurposing drugs provides a new approach to the fight against multidrug-resistant (MDR) bacteria. We have reported that three major tamoxifen metabolites, N-desmethyltamoxifen (DTAM), 4-hydroxytamoxifen (HTAM) and endoxifen (ENDX), presented bactericidal activity against *Acinetobacter baumannii* and *Escherichia coli*. Here, we aimed to analyse the activity of a mixture of the three tamoxifen metabolites against methicillin-resistant *Staphylococcus epidermidis* (MRSE) and *Enterococcus* spp.

MRSE (n=17) and *Enterococcus* spp. (*E. faecalis* n=8, and *E. faecium* n=10) strains were used. MIC of the mixture of DTAM, HTAM and ENDX, and vancomycin were determined by microdilution assay. The bactericidal activity of the three metabolites together and vancomycin against MRSE (SE385 and SE742) and vancomycin-resistant *E. faecalis* (EVR1 and EVR2) strains was determined by time-kill curve assays. Finally, changes in membrane permeability of SE742 and EVR1 strains were analyzed using fluorescence assays.

MIC_50_ and MIC_90_ of tamoxifen metabolites were 1 mg/L for MRSE strains and 2 mg/L for *Enterococcus* spp. strains. In the time-killing assays, tamoxifen metabolites mixture showed bactericidal activity at 2x and 4xMIC for MRSE (SE385 and SE742) and *E. faecalis* (EVR1 and EVR2) strains. This antimicrobial activity of tamoxifen metabolites paralleled an increased membrane permeability of SE385 and EVR2 strains.

Altogether, these results showed that tamoxifen metabolites presented antibacterial activity against MRSE and vancomycin-resistant *E. faecalis*, suggesting that tamoxifen metabolites might increase the arsenal of drugs treatment against these bacterial pathogens.

## Introduction

*Staphylococcus epidermidis* and *Enterococcus* spp. are a common healthcare-associated aetiologies in different infections, causing significant morbidity, mortality and/or healthcare costs (1-3). Glycopeptides are among the recommended treatments for the infections caused by methicillin-resistant *S. epidermidis* (MRSE) and ampicillin-resistant *Enterococcus* spp. (1, 4). However, the emergence of isolates with reduced susceptibility to vancomycin, teicoplanin, linezolid and daptomycin have commonly reported (1, 5-7). Therefore, it is important to increase the arsenal of antimicrobial agents and to find drugs active against MRSE and *Enterococcus* spp. with reduced susceptibility to glycopeptides.

Different approaches can be used to find new antibacterial agents such as repurposing drugs. Anticancer drugs developed to treat breast cancer, like the selective estrogen receptor modulators (SERMs), have been reported to present activity against Gram-positive bacteria (GPB). Clomiphene has demonstrated efficacy against *E. faecium* and *S. aureus* through the inhibition of undecaprenyl diphosphate synthase (UPPS), an enzyme involved in the synthesis of the teichoic acid wall of *S. aureus* (8, 9). Due to this action on the bacterial wall, clomiphene exhibits synergy with β-lactams in restoring methicillin-resistant *S. aureus* susceptibility (9). In addition, tamoxifen was shown to be active against *S. aureus* and *E. faecium in vitro* and in murine and *Galleria mellonella* models of infections, respectively (8, 10). It is noteworthy that tamoxifen antimicrobial activity might also results from cytochrome P450 mediated tamoxifen metabolism releasing three major metabolites N-desmethyltamoxifen (DTAM), 4-hydroxytamoxifen (HTAM) and endoxifen (ENDX) (11). Recently, these three metabolites and their prodrug tamoxifen have been reported to act as a weak base to protect cells and mice against lethal STx1 or STx2 toxicosis (12). Moreover, a previous study from our research group showed that the mixture of DTAM, HTAM and ENDX exhibited MIC_50_ values of 8 and 16 mg/L against clinical isolates of *Acinetobacter baumannii* and *Escherichia coli*, respectively (13); whereas their activity against GPB remain unknown. The objective of this study is to investigate the activity of tamoxifen metabolites against MRSE and *E. faecalis* with reduced susceptibility to vancomycin.

## Results

### Antimicrobial activity of tamoxifen and tamoxifen metabolites

Tamoxifen, tamoxifen metabolites separately and in mixture, and vancomycin were tested against clinical strains of MRSE and *Enterococcus* spp. (*E. faecalis* and *E. faecium*). The MIC_50_ and MIC_90_ values are detailed in Table 1. The MICs of tamoxifen, tamoxifen metabolites mixture and vancomycin for MRSE strains ranged from 2 to 4 mg/L, 0.5 to 2 mg/L and 0.5 to 4 mg/L, respectively; while those for the *Enterococcus* spp. strains ranged from 2 to >32 mg/L, 1 to 4 mg/L and 0.5 to 32 mg/L, respectively. The MIC_50_ and MIC_90_ of tamoxifen for MRSE strains were 2 and 4, respectively, and for *Enterococcus* spp. strains were 4 and 32 mg/L, respectively. The MIC_50_ and MIC_90_ for DTAM, HTAM and ENDX for both pathogens ranged from 2 to 16 mg/L. When these three metabolites were grouped together, their MIC_50_ and MIC_90_ for MRSE and *Enterococcus* spp. were 1 and 2 mg/L, respectively. In the case of vancomycin, the MIC_50_ and MIC_90_ for MRSE strains were 2 and 4 mg/L, respectively, and for *Enterococcus* spp. strains were 1 and 16 mg/L, respectively. These results showed that the mixture of tamoxifen metabolites presented higher antibacterial activity than their prodrug tamoxifen and vancomycin against MRSE and *Enterococcus* spp. strains.

**Table 1.** Minimal inhibitory concentrations effective for ≥50% and ≥90% of isolates tested (MIC_50_ and MIC_90_) of tamoxifen, tamoxifen metabolites and vancomycin for *S. epidermidis* and *Enterococcus* spp.

| Pathogen | N | TAM<br>(mg/L) |  | DTAM<br>(mg/L) |  | HTAM<br>(mg/L) |  | ENDX<br>(mg/L) |  | MET<br>(mg/L) |  | Vancomycin<br>(mg/L) |  |
| --- | --- | --- | --- | --- | --- | --- | --- | --- | --- | --- | --- | --- | --- |
|  |  | MIC<sub>50</sub> | MIC<sub>90</sub> | MIC<sub>50</sub> | MIC<sub>90</sub> | MIC<sub>50</sub> | MIC<sub>90</sub> | MIC<sub>50</sub> | MIC<sub>90</sub> | MIC<sub>50</sub> | MIC<sub>90</sub> | MIC<sub>50</sub> | MIC<sub>90</sub> |
| *S. epidermidis* | 17 | 2 | 4 | 2 | 4 | 8 | 8 | 4 | 8 | **1** | **1** | 2 | 4 |
| *Enterococcus* spp. | 18 | 4 | 32 | 4 | 4 | 8 | 16 | 4 | 8 | **2** | **2** | 1 | 16 |

MET: tamoxifen metabolites, 4-hydroxytamoxifen (HTAM), N-desmethyltamoxifen (DTAM), endoxifen (ENDX) mixture. TAM, tamoxifen; VAN, vancomycin. *Enterococcus* spp., *E. faecalis* and *E. faecium*.

### Time-kill curves

Using time-kill assays, we examined the bactericidal activity of tamoxifen metabolites and vancomycin against MRSE SE385 and SE742 strains and vancomycin-resistant *E. faecalis* EVR1 and EVR2 strains (Figure 1). The MICs of tamoxifen metabolites and vancomycin for these strains are summarized in table 2. Tamoxifen metabolites showed bactericidal activity for both MRSE strains at the highest concentration used (4xMIC), being bactericidal from 2 to 24 h for SE385 strain and at 8 h for SE742 strain; whereas at 2xMIC, tamoxifen metabolites showed bactericidal activity only against SE385 strain at 8 h, but not against SE742 strain. For tamoxifen metabolites at 1xMIC, bacteriostatic activity at any time-point along the assay was observed for SE385 strain and until 8 h for SE742 strain (Figure 1A).

**Table 2.** Minimal inhibitory concentrations tamoxifen metabolites and vancomycin for *S. epidermidis* SE385 an SE742 strains and *E. faecalis* EVR1 and EVR2.

| Strains | MET*<br>(mg/L) | VAN<br>(mg/L) |
| --- | --- | --- |
| *S. epidermidis* SE385 | 1 | 4 |
| *S. epidermidis* SE742 | 1 | 4 |
| *E. faecalis* EVR1 | 2 | 128 |
| *E. faecalis* EVR2 | 1 | 128 |

MET: tamoxifen metabolites 4-hydroxytamoxifen (HTAM), N-desmethyltamoxifen (DTAM), endoxifen (ENDX) mixture.

**Figure 1.**
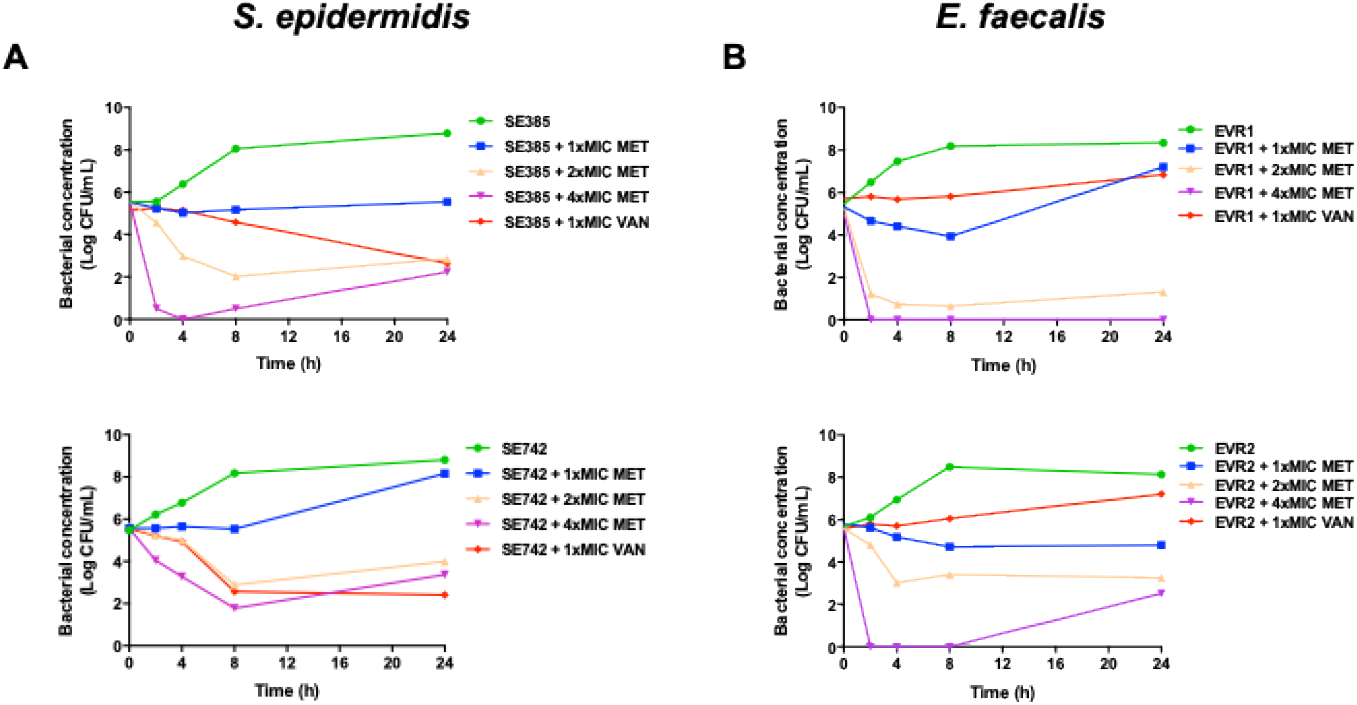
Antibacterial activity of tamoxifen metabolites at different concentrations against *S. epidermidis* and *Enterococcus faecalis* strains. Time-kill curves of *S. epidermidis* SE385 and SE742 strains (**A**) and *E. faecalis* EVR1 and EVR2 strains (**B**) in the presence of 1x, 2x and 4xMIC tamoxifen metabolites and 1xMIC vancomycin for 24 h. MET, tamoxifen metabolites; VAN, vancomycin. Data are represented as mean from two independent experiments.

In the case of vancomycin-resistant *E. faecalis* EVR1 and EVR2 strains, tamoxifen metabolites at 4xMIC showed bactericidal activity for both strains from 2 to 24 h, and at 2xMIC against EVR1. At 1xMIC tamoxifen metabolites, bacteriostatic activity at any time-point along the assay was observed for EVR2 strain and until 8 h for EVR1 strain (Figure 1B). Finally, vancomycin at 1×MIC was bactericidal at 24 h against SE385 and SE742 strains, but not against EVR1 and EVR2 strains.

### Effect of tamoxifen metabolites on the bacterial cell membrane

In order to determine the mode of action of tamoxifen metabolites, we examined their effect on the membrane permeability of SE742 and EVR2 strains. The three tamoxifen metabolites mixture at 0.5xMIC increased significantly the membrane permeability of both strains by 70.22% and 86.6 % (Figure 2). This result suggests that tamoxifen metabolites affect the integrity of the bacterial cell wall of MRSE and vancomycin-resistant *E. faecalis*.

**Figure 2.**
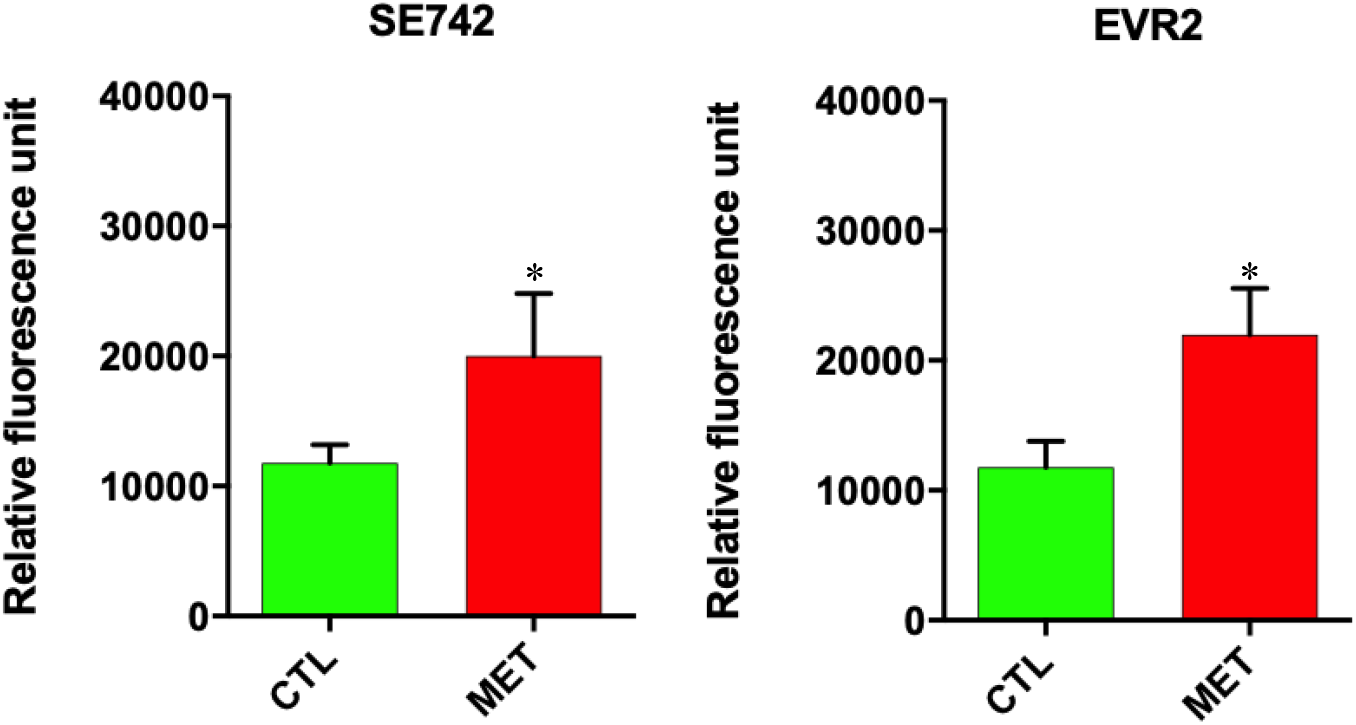
Tamoxifen metabolites effect on the bacterial permeability of *S. epidermidis* and *Enterococcus faecalis* strains. The membrane permeabilization of *S. epidermidis* SE742 and *E. faecalis* VR2 strains in absence and presence of tamoxifen metabolites (0.5xMIC) incubated for 10 min, was quantified by Typhon Scanner. MET: the three tamoxifen metabolites together. CTL: control. * p < 0.05: CTL vs. MET.

## Discussion

The present study provides new data highlighting the antibacterial effect of tamoxifen metabolites against GPB through the increase of bacterial membrane permeability. Among the SERMs investigated, clomiphene and tamoxifen showed activity against *E. faecium* (8, 9). It is worthy of mention that tamoxifen metabolites (HTAM, DTAM and ENDX) showed good activity against *A. baumannii* and *E. coli* (13). We previously showed that tamoxifen metabolites composed of a mixture of HTAM, DTAM and ENDX showed MIC_50_ of 8 and 16 mg/L against 100 and 47 clinical isolates of MDR *A. baumannii* and *E. coli*, respectively (13). Here, the MIC_50_ was 1 and 2 mg/L against MRSE and *Enterococcus* spp., respectively. Compared with MIC_50_ of tamoxifen metabolites against the studied Gram-negative bacteria (GNB), lower MIC_50_ were found against the studied GPB. Obvious reasons for this difference could be due to the structural and molecular differences between the two classes of bacteria (14). Similar differences between GNB and GPB have been observed with other repurposed drugs such as the statin (simvastatin), anthelmintics (niclosamide, oxycloznide and closantel) and anti-inflammatory (celecoxib) drugs (15-17).

It is important to mention that few studies have investigated the activity of tamoxifen metabolites against infectious agents, but there has been no report on their activity against GPB (18). One of these metabolites HTAM has presented activity when used in monotherapy against *Mycobacterium tuberculosis* (MIC_50_ 2.5∼5 mg/L), and in combination with rifampin, isoniazid and ethambutol being the most active at 10 and 20 mg/L of HTAM (19). In addition, HTAM was also reported to be active against *Plasmodium falciparum* and *Cryptococcus neoformans* var. *grubii* (20, 21). HTAM is not the only tamoxifen metabolite used as an infectious agent. The activity of endoxifen was also studied against *C. neoformans* var. *grubii* with MIC of 4 mg/L (21).

Tamoxifen metabolites and vancomycin have different bactericidal activities in time-kill assays. First, at 1xMIC tamoxifen metabolites were not bactericidal against *S. epidermidis*, but vancomycin was bactericidal. Of note, the MIC of vancomycin against MRSE strains is four times higher than the MIC of tamoxifen metabolites. The comparison of the bactericidal activity of both drugs at the vancomycin MIC (4 mg/L) showed higher activity of tamoxifen metabolites than vancomycin. Second, in the case of the *E. faecalis* strains tamoxifen metabolites at 1xMIC were more active than vancomycin at 1xMIC. This activity was higher and faster when both strains of *E. faecalis* were incubated with tamoxifen metabolites at 2x and 4xMIC (4 mg/L) and compared with 4 mg/L vancomycin (Figure 1B). Of note the MIC of tamoxifen metabolites is 64-128 times lower than the MIC of vancomycin.

On the other hand, the bactericidal activity of tamoxifen metabolites against GPB is was higher and faster than that observed against GNB (13). Moreover, tamoxifen metabolites were more bactericidal at lower concentrations against GPB. The fact that tamoxifen metabolites present different activity between GPB and GNB may suggest that the presence of an outer membrane in GNB reduce the activity of tamoxifen metabolites.

In this study, we showed that the three tamoxifen metabolites together produced an increase in membrane permeability of MRSE and vancomycin-resistant *E. faecalis* strains. It is known that the mechanism of action of tamoxifen, the prodrug of DTAM, HTAM and ENDX in fungi is related to the binding to calmodulin (22, 23). Additionally, Scott et al. showed that HTAM might inhibited the phospholipase D in *Pseudomonas aeruginosa* (24). Future studies on the mechanism of action used by tamoxifen metabolites against GPB and on their therapeutic efficacy in animal experimental models of infection would be of interest. In addition to being antibacterial against GPB, tamoxifen and its metabolites have two properties that are advantageous for the treatment of bacterial infections. First, these drugs have excellent bioavailability and, therefore, can be administered orally (25). Second, we have shown that tamoxifen increase the bactericidal activity of macrophages against GNB (26). Probably, this activity may occurred also against GPB.

In conclusion, these results suggest that tamoxifen metabolites are potential antimicrobial agents for use against MRSE and vancomycin-resistant *E. faecalis*, respectively, and it may become, after further development, a possible option for the treatment of infections by MRSE and vancomycin-resistant *Enterococcus* spp.

## Material and methods

### Bacterial strains

Seventeen MRSE and eighteen *Enterococcus* spp. (*E. faecalis* n=8, *E. faecium* n=10) clinical isolates from blood cultures, previously characterized (27, 28) were used in this study. MIC susceptibility breakpoint of vancomycin for both pathogens was determined according to the standard recommendations of the European Committee on Antimicrobial Susceptibility European Committee on Antimicrobial Susceptibility Testing (EUCAST) being susceptible ≤4 mg/L and resistant >4 mg/L (29).

### Antimicrobial agents and reagents

Standard laboratory powders of tamoxifen, HTAM, DTAM, ENDX and vancomycin (Sigma, Spain) were used.

### In vitro susceptibility testing

Minimal inhibitory concentrations (MICs) of HTAM, DTAM and ENDX separately and in mixture, tamoxifen and vancomycin against MRSE and *Enterococcus* spp. strains were determined in two independent experiments by broth microdilution assay according to CLSI guidelines (29). The initial bacterial inoculum 5⨯10^5^ Colony Forming Unit (CFU)/mL for each strain cultured in Mueller Hinton Broth (MHB) (Sigma, Spain) was used in a 96-well plate (Deltlab, Spain) in the presence of HTAM, DTAM, ENDX separately and in mixture (at same concentration), tamoxifen and vancomycin, and incubated for 16-18 h at 37 °C. *E. faecalis* ATCC 29212 and *S. aureus* ATCC 29213 strains were used as control strains. MIC_50_ and MIC_90_, respectively, were determined.

### Time-kill kinetic assays

Time-kill curves of MRSE SE385 and SE742 strains with a vancomycin MIC of 4 mg/L, and *E. faecalis* EVR1 and EVR2 strains with a vancomycin MIC of 128 mg/L were performed in duplicate as previously described (29, 30). Initial inoculums of 5.5⨯10^5^ CFU/mL were added on 5 mL of MHB in presence of 1x, 2x, and 4xMIC of HTAM, DTAM, ENDX mixture, and 1xMIC of vancomycin. Drug free broth was evaluated in parallel as a control. Tubes of each condition were incubated at 37 °C with shaking (180 rpm), and viable counts were determined by serial dilution at 0, 2, 4, 8, and 24 h. Viable counts were determined by plating 100 μL of control, test cultures, or the respective dilutions at the indicated times onto sheep blood agar plates (ThermoFisher, Spain). Plates were incubated for 24 h at 37 °C, and, after colony counts, the log_10_ of viable cells (CFU/mL) was determined. Bactericidal was defined as a reduction of ≥3 log_10_ CFU/mL with the initial inoculum.

### Membrane permeability Assays

Bacterial suspensions (adjusted to optical density at 600 nm = 0.2) of SE742 and VR2 strains were placed on a 96-well plate, incubated with 0.5xMIC of tamoxifen metabolites mixture and mixed in a solution of phosphate buffered saline containing Ethidium Homodimer-1 (EthD-1) (1:500) (Invitrogen, Carlsbad, CA, USA). After 10 min of incubation, fluorescence was monitored during 160 min using a Typhoon FLA 9000 laser scanner (GE Healthcare Life Sciences, Marlborough, MA, USA) and quantified with ImageQuant TL software (GE Healthcare Life Sciences, USA). Bacterial counts were obtained at the beginning and end of the experiment to ensure that the metabolite mixture did not present bactericidal activity against *S. epidermidis* and *Enterococcus* spp. strains.

### Statistical analysis

Group data were presented as means ± standard errors of means (SEM). Difference in membrane permeability were assessed by Student t-test. The SPSS (version 23.0; SPSS Inc., Armonk, NY, USA) statistical package was used.

## Acknowledgments

We thank Dr. Diego Garcia Martínez de Artola for the kind gift of the *E. faecalis* EVR1 and EVR2 strains.

## Funding

This study was supported by the Instituto de Salud Carlos III, Proyectos de Investigación en Salud (grants CP15/00132, PI16/01378 and PI19/01453) and by Plan Nacional de I+D+i 2013-2016 and Instituto de Salud Carlos III, Subdirección General de Redes y Centros de Investigación Cooperativa, Ministerio de Ciencia, Innovación y Universidades, Spanish Network for Research in Infectious Diseases (REIPI RD16/0016/0009) -co-financed by European Development Regional Fund “A way to achieve Europe”, Operative program Intelligent Growth 2014-2020. Younes Smani is supported by the Subprograma Miguel Servet Tipo I from the Ministerio de Economía y Competitividad of Spain (CP15/00132).

## Author contributions

Y.S. conceived the study and designed the experiments. A.M.C., A.V.D., R.A.A, performed experiments and interpreted data. M.E.J.M., J.P. revised the manuscript and Y.S. wrote the manuscript with the input of all the other authors.

## Competing interests

No conflicts of interest to declare.

## References

1. Becker K, Heilmann C, Peters G. 2014. Coagulase-negative staphylococci. Clin Microbiol Rev 27:870–926.

2. Heilmann C, Ziebuhr W, Becker K. 2019. Are coagulase-negative staphylococci virulent? Clin Microbiol Infect 25:1071–1080.

3. Chiang HY, Perencevich EN, Nair R, Nelson RE, Samore M, Khader K, Chorazy ML, Herwaldt LA, Blevins A, Ward MA, Schweizer ML. 2017. Incidence and outcomes associated with infections caused by vancomycin-resistant Enterococci in the United States: systematic literature review and meta-analysis. Infect Control Hosp Epidemiol 38:203–215.

4. Girón-González JA, Pérez-Cano R. 2003. Tratamiento de las infecciones por enterococo. Rev Clin Esp 203:482–485.

5. Pfaller MA, Sader HS, Flamm RK, Castanheira M, Smart JI, Mendes RE. 2017. In vitro activity of telavancin against clinically important Gram-positive pathogens from 69 U.S. medical centers (2015): Potency analysis by U.S. census divisions. Microb Drug Resist 23:718–726.

6. Santerre Henriksen A, Smart J, Hamed K. 2018. Comparative activity of ceftobiprole against coagulase-negative staphylococci from BSAC bacteraemia surveillance programme 2013-2015. Eur J Clin Microbiol Infect Dis 37:1653–1659.

7. Levitus M, Rewane A, Perera TB. Vancomycin-Resistant Enterococci (VRE). In: StatPearls [Internet]. Treasure Island (FL): StatPearls Publishing; 2020 Jan–.2020 Mar 30.

8. Jacobs AC, Didone L, Jobson J, Sofia MK, Krysan D, Dunman PM. 2013. Adenylate kinase release as a high-throughput-screening-compatible reporter of bacterial lysis for identification of antibacterial agents. Antimicrob Agents Chemother 57:26–36

9. Farha MA., Czarny TL, Myers CL, Worrall LJ, French S, Conrady DG, Wang Y, Oldfield E, Strynadka NCJ, Brown ED. 2015. Antagonism screen for inhibitors of bacterial cell wall biogenesis uncovers an inhibitor of undecaprenyl diphosphate synthase. Proc Natl Acad Sci USA 112:11048–11053.

10. Corriden R, Hollands A, Olson J, Derieux J, Lopez J, Chang JT, Gonzalez DJ, Nizet V. 2015. Tamoxifen augments the innate immune function of neutrophils through modulation of intracellular ceramide. Nat Commun 6:8369.

11. Klein DJ, Thorn CF, Desta Z, Flockhart DA, Altman RB, Kleina TE. 2013. PharmGKB summary: tamoxifen pathway, pharmacokinetics. Pharmcogenet Genomics 23:643–647.

12. Selyunin AS, Hutchens S, McHardy SF, Mukhopadhyay S. 2019. Tamoxifen blocks retrograde trafficking of Shiga toxin 1 and 2 and protects against lethal toxicosis. Life Sci Alliance 2:e201900439.

13. Miró-Canturri A, Ayerbe-Algaba R, Vila-Domínguez A, Jiménez-Mejías ME, Pachón J, Smani Y. 2021. Repurposing of the tamoxifen metabolites to combat infections by multidrug-resistant Gram-negative bacilli. Antibiotics 10:336.

14. Brown L, Wolf JM, Prados-Rosales R, Casadevall A. 2015. Through the wall: extracellular vesicles in Gram-positive bacteria, mycobacteria and fungi. Nat Rev Microbiol 13: 620–630.

15. Thangamani S, Mohammad H, Abushahba MF, Hamed MI, Sobreira TJ, Hedrick VE, Paul LN, Seleem MN. 2015. Exploring simvastatin, an antihyperlipidemic drug, as a potential topical antibacterial agent. Sci Rep 5:16407.

16. Rajamuthiah R, Fuchs BB, Conery AL, Kim W, Jayamani E, Kwon B, Ausubel FM, Mylonakis E. 2015. Repurposing salicylanilide anthelmintic drugs to combat drug resistant Staphylococcus aureus. PLoS One 10:e0124595.

17. Thangamani S, Younis W, Seleem MN. 2015. Repurposing celecoxib as a topical antimicrobial agent. Front Microbiol 6:750.

18. Montoya MC, Krysan DJ. 2018. Repurposing estrogen receptor antagonists for the treatment of infectious disease. mBio 9:e02272–18.

19. Chen FC, Liao YC, Huang JM, Lin CH, Chen YY, Dou HY, Hsiung CA. 2014. Pros and cons of the tuberculosis drugome approach-an empirical analysis. PLoS One 9:e100829.

20. Weinstock A, Gallego-Delgado J, Gomes C, Sherman J, Nikain C, Gonzalez S, Fisher E, Rodriguez A. 2019. Tamoxifen activity against Plasmodium in vitro and in mice. Malar J 18:378.

21. Butts A, Koselny K, Chabrier-Rosello Y, Semighini CP, Brown JC, Wang X, Annadurai S, DiDone L, Tabroff J, Childers WE, J., Abou-Gharbia M, Wellington M, Cardenas ME, Madhani HD, Heitman J, Krysan DJ. 2014. Estrogen receptor antagonists are anti-cryptococcal agents that directly bind EF hand proteins and synergize with fluconazole in vivo. mBio 5:e00765–13.

22. Dolan K, Montgomery S, Buchheit B, DiDone L, Wellington M, Krysan DJ. 2009. Antifungal activity of tamoxifen: in vitro and in vivo activities and mechanistic characterization. Antimicrob Agents Chemother 53:3337–3346.

23. Butts A, Martin JA, DiDone L, Bradley EK, Mutz M, Krysan J. 2015. Structure-activity relationships for the antifungal activity of selective estrogen receptor antagonists related to tamoxifen. Plos ONE 10:e0125927.

24. Scott SA, Spencer CT, O’Reilly MC, Browb KA, Lavieri RR, Cho CH, Jung DI, Larock RC, Brown HA, Lindsley CW. 2015. Discovery of desketoraloxifene analogues as inhibitors of mammalian, Pseudomonas aeruginosa, and nape phospholipase d enzymes. ACS Chem Biol 10:421–432.

25. Morello KC, Wurz GT, DeGregorio MW. 2003. Pharmacokinetics of selective estrogen receptor modulators. Clin Pharmacokinet 42:361–372.

26. Miró-Canturri A, Ayerbe-Algaba R, del Toro R, Pachón J, Smani Y. 2020. Tamoxifen repurposing to combat infections by multidrug-resistant Gram-negative bacilli. bioRxiv. doi: https://doi.org/10.1101/2020.03.30.017475.

27. Domínguez-Herrera J, Docobo-Pérez F, López-Rojas R, Pichardo C, Ruiz-Valderas R, Lepe JA, Pachón J. 2012. Efficacy of daptomycin versus vancomycin in an experimental model of foreign-body and systemic infection caused by biofilm producers and methicillin-resistant Staphylococcus epidermidis. Antimicrob Agents Chemother 56:613–617.

28. Vila Domínguez A, Ayerbe-Algaba R, Miró Canturri A, Rodríguez-Villodres A, Smani Y. 2020. Antibacterial activity of colloidal silver against Gram-negative and Gram-positive bacteria. Antibiotics 9:36.

29. Clinical and Laboratory Standards Institute. Performance Standards for Antimicrobial Susceptibility Testing. 29^th^ ed. CLSI supplement M100. CLSI: Wayne, PA, USA, 2019.

30. Miró-Canturri A, Ayerbe-Algaba R, Villodres ÁR, Pachón J, Smani Y. 2020. Repositioning rafoxanide to treat Gram-negative bacilli infections. J Antimicrob Chemother 75:1895–1905.

